# Differential adaptive potential and vulnerability to climate-driven habitat loss in Brazilian mangroves

**DOI:** 10.1101/2022.02.11.480143

**Authors:** João de Deus Vidal, Gustavo Maruyama Mori, Mariana Vargas Cruz, Michele Fernandes da Silva, Yohans Alves de Moura, Anete Pereira de Souza

## Abstract

Geographic and environmental differences have been identified as factors influencing Brazilian mangrove trees’ genetic diversity. Geographically, distinct species have convergent spatial genetic structures, indicating a limited gene flow between northern and southern populations. Environmentally, genomic studies and common garden experiments have found evidence of local adaptations along the latitudinal gradient of the Brazilian coast. However, little is known about how such adaptive heterogeneity could be affected by a rapidly changing climate in the coming decades, and the combination of deforestation and climate-induced habitat loss may affect these forests and their genetic diversity. Here, we applied two genomic-environmental association methods to model the turnover of potentially adaptive alleles for two dominant mangrove trees: *Avicennia germinans* and *A. schaueriana*. We analyzed a total of 134 individuals from six populations of *A. germinans* and ten populations of *A. schaueriana* spanning the Brazilian coast from 1 °S to 28 °S. Gradient forest models identified temperature-related variables as the most important predictors for *A. germinans* outlier loci, whereas both temperature and precipitation were important for *A. schaueriana*. We modeled allele frequencies and projected them for future climatic scenarios to estimate adaptively driven vulnerability. We assessed climate-driven habitat loss through climate-only distribution models and calculated annual deforestation rates for each sampled region. Finally, to assess the vulnerability of individual populations, we combined the environmental suitability, deforestation data, and adaptive vulnerability projections. For both species, subtropical populations presented a higher vulnerability than equatorial populations to climate-driven habitat loss. We also identified deforestation rates at the sampled sites that were alarmingly higher than the global average mangrove deforestation rate. Our results provide improved estimates of the impacts of ongoing climate change and human-caused habitat loss on the distribution of mangroves and highlight the importance of site-based conservation strategies that consider individual subtropical and equatorial mangrove forests.

## 1. Introduction

The potential of species to respond to the rapid pace of human-induced climatic change constitutes a major concern for biological conservation worldwide. Model-based estimates indicate that under a moderate carbon dioxide emission scenario, almost 25% of species across most taxonomic groups could face extinction by 2050 (Thomas et al. 2004). Communities in coastal environments, such as mangrove forests, are among the most vulnerable systems due to the high specificity of their niches, the sea-level rise predicted for this century (Gilman et al. 2008, Sippo et al. 2018, Friess et al. 2018), and the pace of climate change (Loarie et al. 2009). Given the ecological relevance of these communities for carbon fixation (Eong 1993) and habitat formation (Tomlinson 1986) and their high vulnerability (Hoegh-Guldberg & Bruno 2010, Polidoro et al. 2010), mangroves constitute a key target for biodiversity conservation and climate change mitigation.

The strong human pressure on coastal regions and the historically high deforestation rates in mangroves constitute additional challenges to the long-term viability of mangrove populations under future climatic scenarios. These factors negatively impact the availability and connectivity of habitats, compromising the ability of mangrove species to occupy areas within their climatic tolerances (Jump & Peñuelas 2005). However, despite the observed progress in mangrove conservation during the last decade (Goldberg et al. 2020), these forests are still declining globally by approximately 0.4% of their area per year and are considered critically endangered in 26 out of the more than 100 countries where they occur (Duke et al. 2007).

Although human-driven deforestation is the main threat to mangroves, habitat loss may also affect the long-term capacity of their populations to persist and respond to future climatic scenarios. Ongoing climatic changes are promoting range shifts, especially toward higher latitudes, for several species of animals and plants (Chen et al. 2011), including mangroves (Osland et al. 2016). However, this shift depends on the survivability, dispersal capacity, and migration rates of the species, factors that were limited in plants during past climatic oscillations (Davis & Shaw 2001). Additionally, climate change has been linked to higher mortality and other negative effects on mangrove forests (Duke et al. 2017, Lovelock et al. 2017), although these effects may be highly variable between populations along environmental gradients.

In South America, mangroves occupy ca. 2 million hectares, mostly along the Brazilian coast (FAO 2007), with estimated impacts on the regional economy between US $33,000 and 57,000 per hectare per year (UNEP, 2014). In Brazil, due to the country’s wide geographic extent that ranges from 33.75°S to 5.27°N and the variety of environmental conditions found along the coastal latitudinal gradient, mangrove forests are naturally exposed to widely variable adaptive pressures (Cruz et al. 2020, da Silva et al. 2021). While populations from the northern coast of the country inhabit equatorial environments with a high annual rainfall and warmer temperatures, southern populations reach latitudes as low as 28°S and occupy subtropical areas with lower temperatures and less annual precipitation. This climatic variability has been associated with differentiation in the adaptive genetic composition of the mangrove populations, especially in loci related to temperature, solar radiation, and water stress (Bajay et al. 2018, Cruz et al. 2020, da Silva et al. 2021). As a result, temperature rise and changes in precipitation patterns may differentially affect the persistence and survival of populations due to local adaptations, such as those reported for the dominant species *Avicennia germinans* and *A. schaueriana* (*Acanthaceae*) (Cruz et al. 2020, da Silva et al. 2021).

Combined with environmentally driven divergence, neutral forces play a key role in population genetics. Mangrove trees have high dispersal potential due to their waterborne propagules (fruits or seeds), but long-distance dispersal events are rare, which causes limited connectivity among populations (Van der Stocken et al. 2019). At regional and biogeographic scales, abiotic factors such as coastal geomorphology (e.g., Triest et al. 2020), landmasses (e.g., Cerón-Souza et al. 2015, Ochoa-Zavala et al. 2019, Mori et al. 2021), and ocean currents and eddies (e.g., Mori et al. 2015, Cerón-Souza et al. 2015, Kennedy et al. 2016, Triest et al. 2021, da Silva et al. 2021) influence how propagules move across geographical space. As these factors shape propagule dispersal, they may lead to the differentiation of populations and, consequently, their genetic divergence. Restricted gene flow, in turn, may facilitate local adaptation in each population (Kawecki & Ebert 2004, Savolainen et al. 2007). Despite advances in scientific knowledge on population genetics, for most species, especially those in South American mangroves (Cruz et al. 2019), little is known about the distribution of adaptive variation, making it difficult to estimate the capacity of these plants to adapt to novel climates and to develop suitable conservation efforts.

Recently, methodological advances have made it possible to model how differences in adaptation along environmental gradients may affect the vulnerability of individual populations to climatic changes (Fitzpatrick & Keller 2015, Bay et al. 2018). These approaches take into account the frequencies of potentially adaptive alleles identified with environmental-genotypic correlation methods and apply statistical models to simulate the required allele frequencies under future environmental projections. Combined with common garden experiments, this type of simulation may provide powerful insights into the adaptive capacity of populations under environmental changes (Fitzpatrick et al. 2021). Here, we implemented a genomic-environmental approach to predict how populations of two mangrove species, *A. germinans* and *A. schaueriana*, may adapt to future climatic conditions. We tested the association between adaptive allelic diversity and modern climatic conditions and projected populations’ allelic diversity into the geographic space under future climatic scenarios. We estimated individual populations’ risks of both climate-induced habitat loss and deforestation and discussed the implications of these data on the long-term viability of these forests. Our findings highlight the need to develop specific population-focused conservation strategies and highlight the importance of considering local adaptations in species conservation.

## Materials and Methods

### Population sampling and genomic sequencing

The overall methodology is described in Figure 1. We used a genomic dataset previously obtained by our research group (Cruz et al. 2019, 2020) via high-throughput DNA sequencing to model the adaptive potential of *A. germinans* and *A. schaueriana* populations under future climate scenarios. The dataset comprises 57 adult *A. germinans* trees sampled at six populations spanning latitudes from −0.71° to −8.59° and 77 adult *A. schaueriana* trees sampled at ten populations spanning latitudes from −0.82° to −28.48° (Fig. 2, Table 1). Full population information is available in Table 1. For each individual, total DNA was previously extracted using the DNeasy Plant Mini Kit (Qiagen, Hilden, Germany) and NucleoSpin Plant II Kit (Macherey Nagel). We used NEXTera-tagmented, reductively amplified DNA (nextRAD) library preparation, a method that uses Nextera technology (Illumina Inc., USA) to simultaneously fragment and tag target DNA with sequencing adapters (Russello et al. 2015) to identify and genotype SNPs. The methods used to build and sequence nextRAD libraries and to process reads are described in the original manuscripts (Cruz et al. 2019, 2020). The sequences obtained through the nextRAD libraries were filtered by Cruz et al. (2020) and Cruz et al. (2019) using a maximum threshold of 65% for missing data, Phred scores greater than 30, with 8× minimum coverage, a single SNP per locus, and a minor allele frequency ≥0.05 with vcftools v.0.1.12b46 (Danecek et al., 2011). The dataset we used comprised 2,297 and 6,170 SNP loci for *A. germinans* and *A. schaueriana*, respectively, which were then used in the genome-environment association step.

**Fig. 1.**
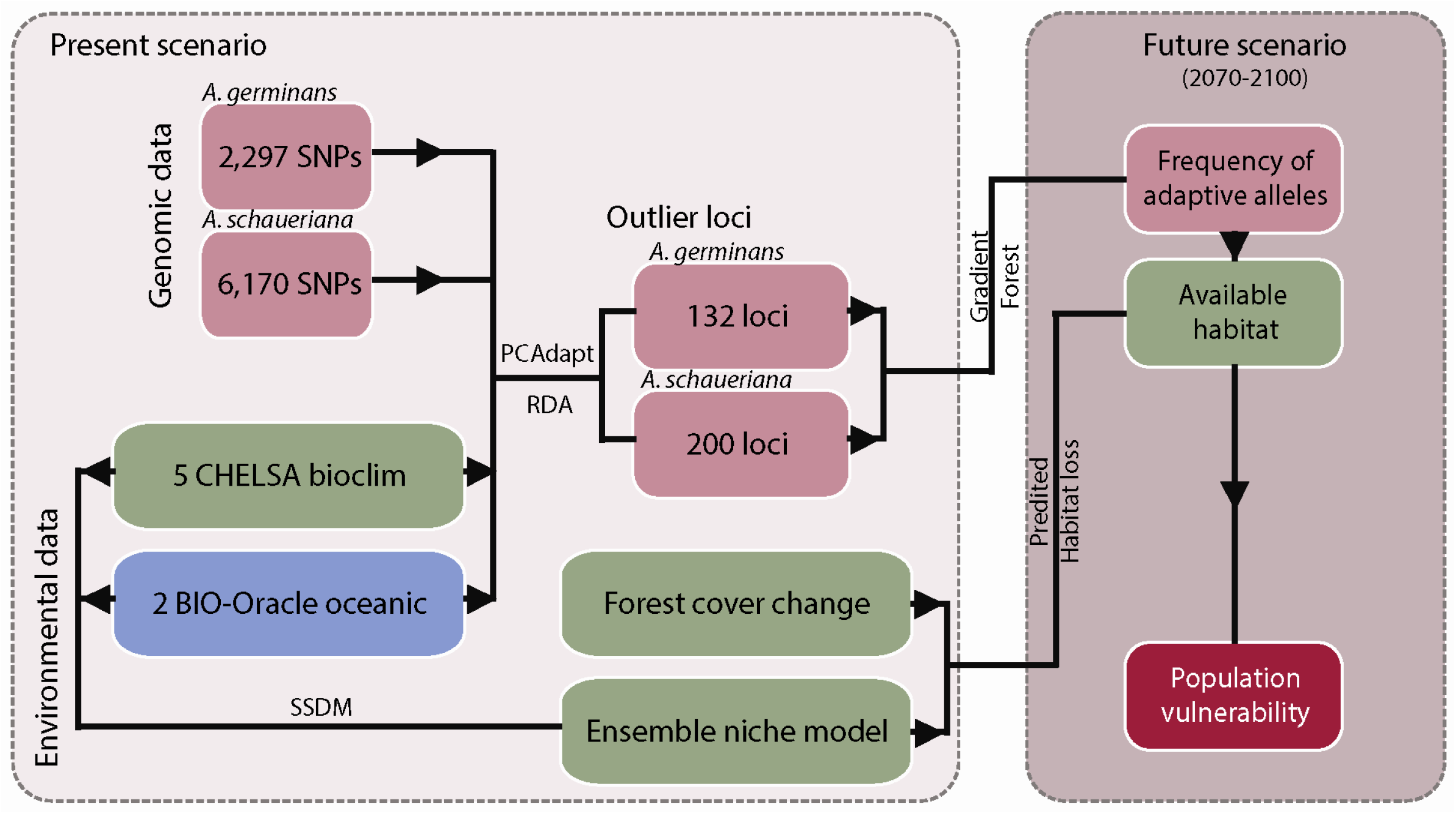
Diagram of the analyses and datasets applied to estimate the vulnerability of individual populations to climate change based on the projected adaptive allelic frequencies and habitat loss.

**Fig. 2.**
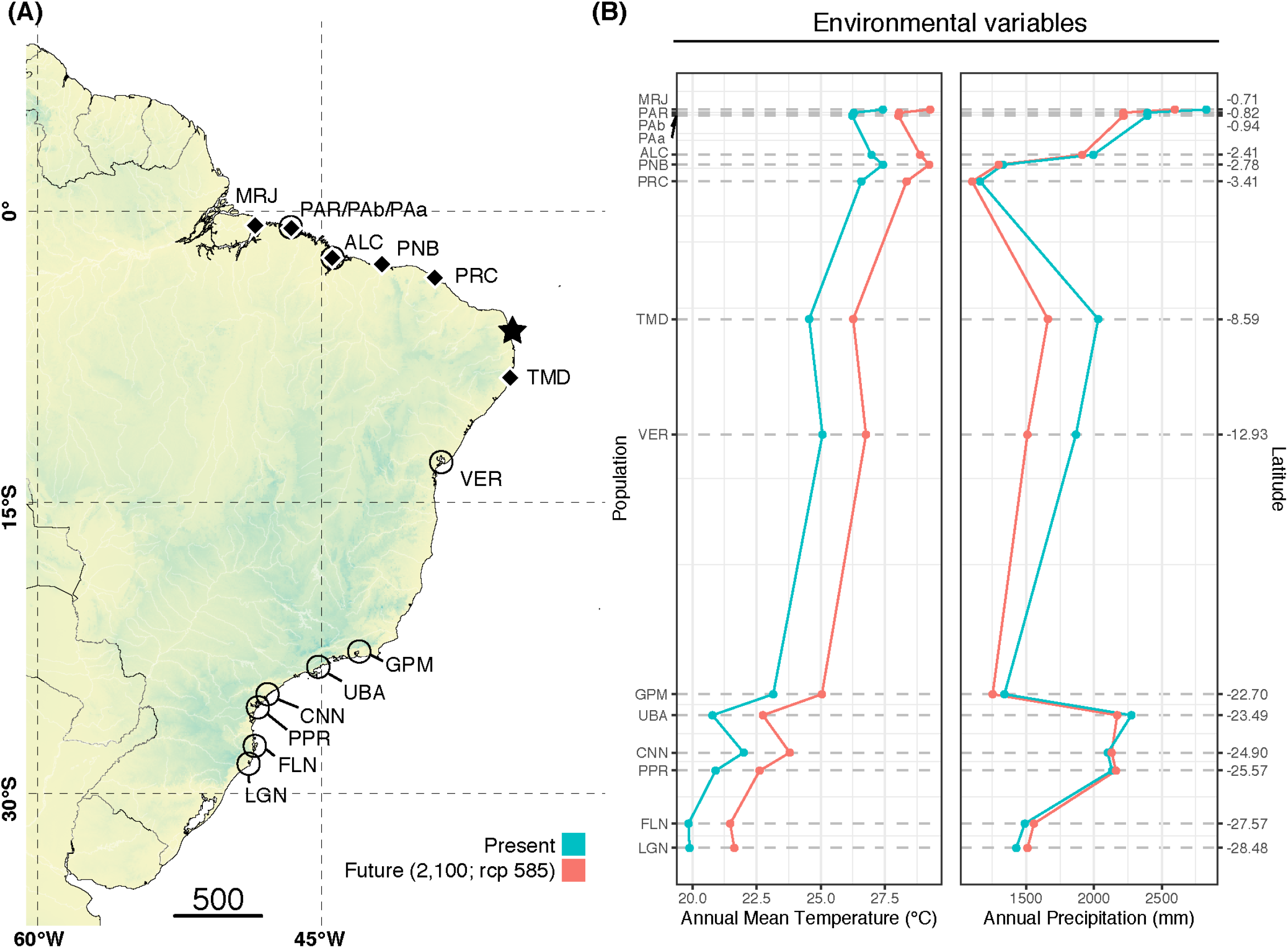
**(A)** Sampled locations for *Avicennia germinans* (black diamonds) and *Avicennia schaueriana* (empty circles) and the northeastern extremity of South America (NEESA) (star). **(B)** Present and future (2100; SSP5-8.5) climatic conditions for each site based on WorldClim 2.0 (Fick & Hijmans 2017).

**Table 1.** Population name codes, geographic coordinates, and sample sizes of *Avicennia germinans* and *Avicennia schaueriana* individuals sampled for the genomic environmental association analysis.

| Species | Population | N | Latitude | Longitude |
| --- | --- | --- | --- | --- |
| *Avicennia germinans* | MRJ | 8 | -0.70565 | -48.48630 |
|  | PAB | 18 | -0.93917 | -46.72139 |
|  | ALC | 5 | -2.40971 | -44.40573 |
|  | PNB | 9 | -2.78051 | -41.82358 |
|  | PRC | 7 | -3.41269 | -39.05708 |
|  | TMD | 10 | -8.58974 | -35.06445 |
| *Avicennia schaueriana* | PAR | 9 | -0.82377 | -46.61650 |
|  | ALC | 6 | -2.40971 | -44.40573 |
|  | PRC | 9 | -3.41269 | -39.05708 |
|  | VER | 9 | -12.93400 | -38.67420 |
|  | GPM | 9 | -22.69890 | -43.00152 |
|  | UBA | 9 | -23.49000 | -45.16300 |
|  | CNN | 8 | -24.89710 | -47.84720 |
|  | PPR | 8 | -25.57500 | -48.35250 |
|  | FLN | 7 | -27.56780 | -48.51890 |
|  | LGN | 3 | -28.48460 | -48.84240 |

**Table 2.** Overall importance of variables according to the impurity reduction measured by the Gini index (Breiman et al., 1984) for *Avicennia germinans* and *Avicennia schaueriana* outlier and reference loci (i.e., neutral) calculated by the gradient forest models.

| Environmental predictor | *Avicennia germinans* |  | *Avicennia schaueriana* |  |
| --- | --- | --- | --- | --- |
|  | Overall importance (reference) | Overall importance (outlier) | Overall importance (reference) | Overall importance (outlier) |
| Mean annual air temperature | 0.02 | **0.11** | 0.07 | **0.17** |
| Annual precipitation amount | 0.05 | 0.02 | 0.01 | 0.05 |
| Precipitation seasonality | 0.04 | **0.11** | 0.07 | **0.16** |
| Isothermality | 0.04 | **0.08** | 0.06 | **0.15** |
| Annual range of air temperature | 0.04 | **0.13** | 0.07 | **0.19** |
| Distance | 0.04 | **0.09** | 0.04 | **0.08** |
| Growing season length | 0.04 | 0.02 | 0.02 | 0.04 |
| Ocean salinity | 0.04 | **0.09** | 0.02 | 0.1 |
| Ocean surface temperature | 0.02 | 0.01 | 0.06 | **0.11** |

**Table 3.**
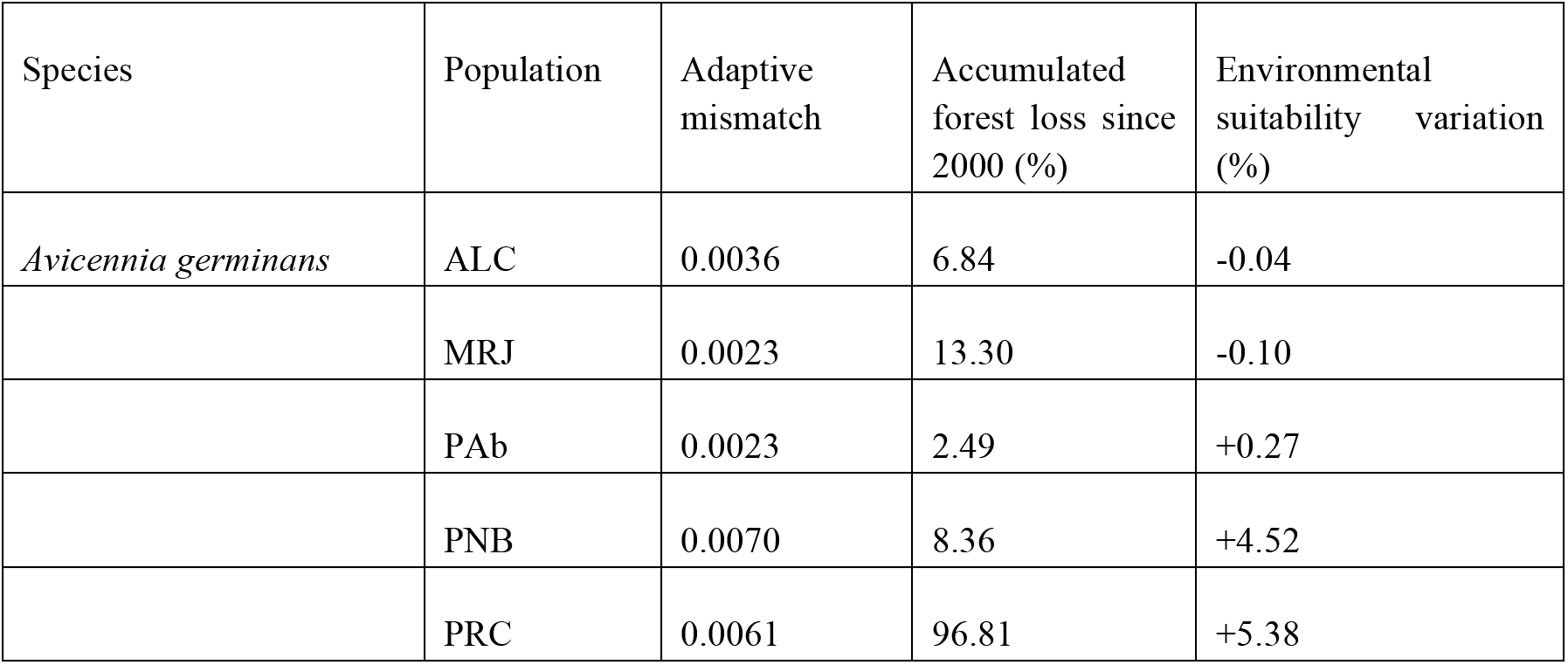
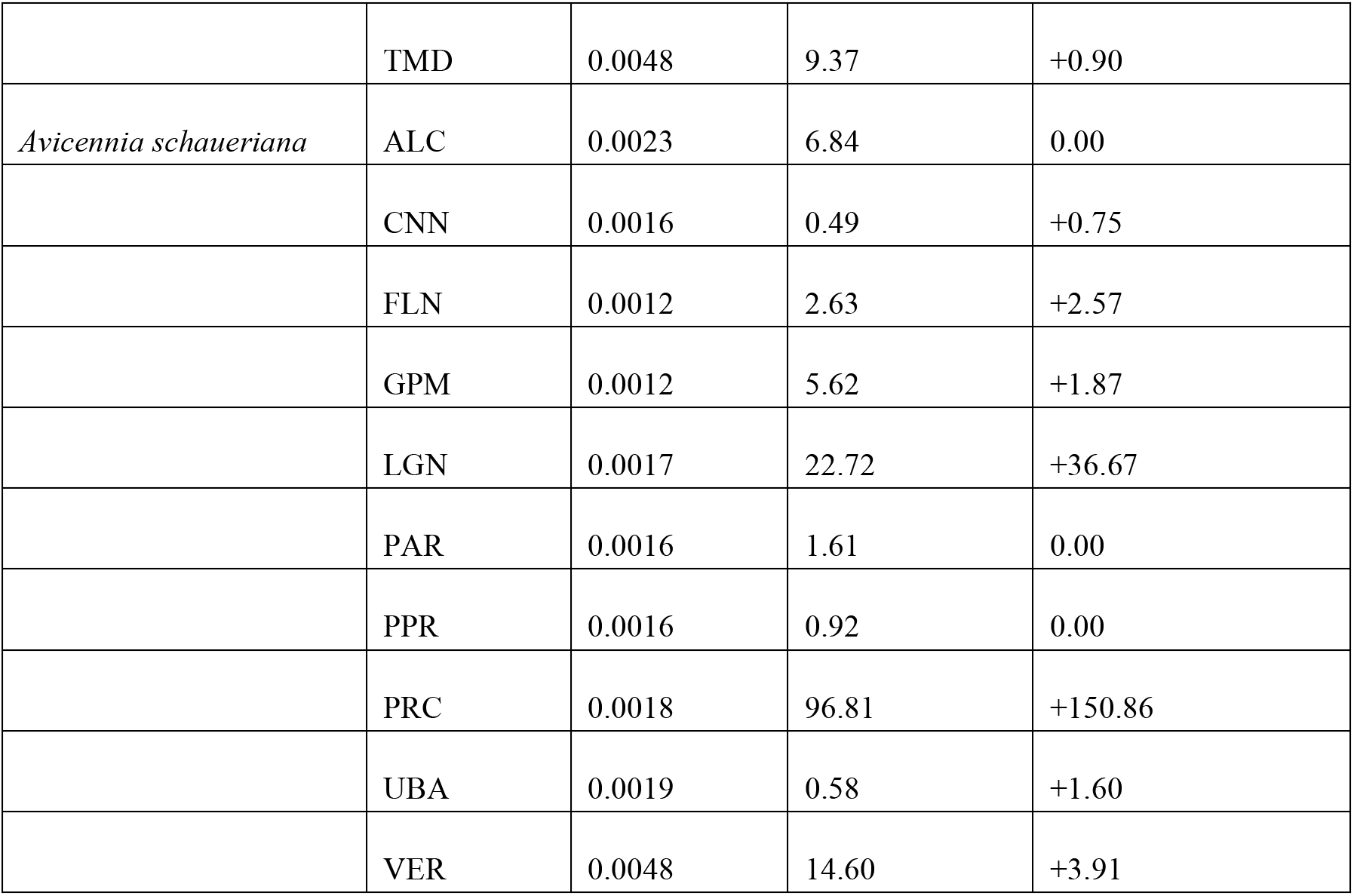
Summarized variations in present and future (2100, RCP85/SSP-85) adaptive allelic frequencies (Procrustes differences between present and future scenarios), accumulated forest loss since 2000 (%) based on current local deforestation rates, and suitability differences based on ensemble presence-absence species distribution models (%).

### Genomic scan for loci under adaptation and allelic frequency modeling

We followed the method by Fitzpatrick & Keller (2015), who developed an approach for adapting a community turnover modeling method based on the gradient forest (GF) algorithm (Smith & Ellis, 2020) applied to allelic frequency datasets along environmental gradients. The GF algorithm applies a machine-learning algorithm to subset values of allele frequencies and associates these values with transitions along gradients of environmental variables. By doing so, it is possible to evaluate the biological variation across environmental gradients and to project that variation to future climatic scenarios (Fitzpatrick & Keller 2015). In the first step, we identified loci that are potentially under selection by scanning genome-wide datasets for outlier loci using the package “PCAdapt” version 4.3.3 (Luu et al. 2020) for the R platform (R core team, 2018); this package detects loci-correlated population structures using a false discovery rate (FDR) of <0.1. Using environmental and geographic distance variables as predictors, this test can calculate the z scores obtained when adjusting SNPs to the selected principal components, providing a measurement of the deviation of each locus from the mean genetic variation.

To minimize the occurrence of false positives, in addition to using PCAdapt, we also implemented an additional genotype-environment association (GEA) method based on redundancy analysis (RDA) developed by Forester et al. (2018). RDA is also based on ordination, but unlike PCAdapt, RDA is able to constrain the analysis with environmental variables and is more efficient at detecting true positives under certain evolutionary scenarios. We ran RDA using the R package “vegan” version 2.5-7 (Oksanen et al., 2020), using a twofold standard deviation cutoff (p < 0.05) as a threshold. We overlaid both lists of candidate loci and excluded loci that were not simultaneously identified by both methods.

We subset the genomic dataset into a reference dataset with 300 randomly selected reference SNPs and an outlier dataset comprising all outlier SNPs. For each locus, we calculated individual population allelic frequencies using the function “makefreq()” from the package adegenet version 2.1.5 (Jombart 2008). We fitted gradient forest models for both the reference and outlier SNPs vs. environmental predictors using the function “gradientForest()” in the R package gradientForest version 0.1-18 (Smith & Ellis, 2020) using 500 bootstrapped trees and no transformation of the dataset. We calculated the importance of each environmental predictor in the allelic frequency of both species by using the impurity reduction measured by the Gini index (Breiman et al., 1984). To measure the vulnerability to future climatic conditions, we compared the spatial patterns of potential adaptive genetic variation between current and future climates with a modified Procrustes analysis using the principal components computed from PCAdapt, following Fitzpatrick & Keller (2015). Procrustes analysis compares matrices in a dataset by “rotating a matrix to maximum similarity with a target matrix, minimizing the sum of squared differences” (from Oksanen et al. 2020). This approach has been applied to compare ordination results in genomic modeling studies (Fitzpatrick & Keller, 2014) and is particularly useful for comparisons using multidimensional scaling (Oksanen et al. 2020). The implementation of the method to raster objects in R is based on the function “procrustes()” of the package “vegan”, as applied in Fitzpatrick & Keller (2015) and modified by Maier (2018).

### Environmental analysis and distribution modeling

We used models generated with the GF algorithm to estimate the geographic distributions of allelic frequencies and to project them to future climate scenarios. To generate the models and identify PCA-significant loci, we used the set of 30-arc second resolution bioclimatic variables from CHELSA version 2.1 (Karger et al. 2017) and Bio-ORACLE version 2.2 mean surface temperature and salinity data (Assis et al. 2018). To minimize overfitting in the modeling steps, we performed a variable selection step by randomly sampling 1,000 points within the region of interest using the function “sampleRandom()” in the R package raster version 3.4–13 (Hijmans, 2021) and calculating Pearson’s correlation for the stack of eight environmental predictors (two oceanic and five bioclimatic) using the R function “cor()” from the R package stats version 3.6.3 (R Core Team, 2020). We randomly removed variables from highly correlated pairs (i.e., Pearson’s correlation > 0.7) and restricted the environmental dataset to include only the less correlated environmental predictors: oceanic salinity and oceanic surface temperature, bio1 (mean annual air temperature), bio3 (isothermality), bio7 (annual range of air temperature), bio12 (annual precipitation amount), bio15 (precipitation seasonality), and gsl (growing season length). We also obtained the geographic distance between pairs of populations from Silva et al. (2020), which calculated the pairwise distances based on the coastline extent. For the future scenario, we used projected datasets for the years 2070-2100 for the bioclimatic predictors under the SSP5-8.5 emission scenario based on the Max Planck Institute Earth System Model (MPI-ESM1.2) (Gutjahr et al., 2019), which combines high-resolution circulation models for the atmospheric and oceanic climate mean states. For the Bio-ORACLE dataset, the future scenario predictors available are projected for the years 2090-2100 based on CMIP5 RCP 85, which is the previous version of SSP5-8.5. Future climatic projections are developed and released by the World Climate Research Programme (WCRP) Coupled Model Intercomparison Project (CMIP). Climate models featured in CMIP include one “very high baseline scenario”, namely, RCP 85 on CMIP5 (Taylor et al. 2012) and SSP5-8.5 on CMIP6 (Eyring et al., 2016). Both models depict the highest emissions, no-policy baseline scenario, with SSP5-8.5 showing approximately 20% higher CO2 emissions by 2100 and lower emissions of other greenhouse gases than its previous version, the RCP8.5 scenario, which estimates a 3.7 °C (ranging from 2.6 to 4.8) increase in temperature by 2100. We generated a 50 km buffer around each occurrence point and extracted the values of the variables within the area. Next, we used a thin-plate spline regression function from the “fields” package version 12.5 (Nychka et al. 2017) to interpolate values for continental-scale cells—a necessary step for the projection of the generated distribution models. We also obtained the geographic distance between pairs of populations from Silva et al. (2020), which calculated the pairwise distances based on the coastline extent. For the modeling step, we adopted the nearest population distance as a measure of the relative geographic isolation of each population.

To compare the predictions of genomic and nongenomic models, we also generated and projected ensemble distribution models based on only environmental factors using the R package “SSDM” version 0.2.8 (Schmitt et al. 2017). We downloaded and filtered records for both species from the Global Biodiversity Information Facility (GBIF) using the R package coordinateCleaner version 2.0–18 (Zizka et al. 2019) and obtained a total of 1,350 occurrence records for *A. germinans* and 373 for *A. schaueriana*. We modeled each species distribution, combining generalized linear model (GLM), artificial neural network (ANN), support vector machine (SVM), multivariate adaptive regression spline (MARS), and random forest (RF) methods, aviilable in SSDM, which were demonstrated to show similar high sensitivity–specificity ratios for narrow-niche species (Qiao et al. 2015). By adopting an ensemble modeling approach, we aimed to account for the differential performances of modeling algorithms while ensuring that the results reflected the most accurate projections. One repetition per algorithm and the kappa value were used as the model-weighted ensemble metrics for evaluation. To calculate the importance of each environmental predictor in the geographic distribution of both species, we calculated Pearson’s correlation r between the predictions of each model and a model calculated after removing that variable to obtain a value of *1-r*, indicating the relative importance of each predictor to the model. We also projected the ensemble models to the SSP5-8.5 scenario. Finally, we converted the present and predicted distributions into presence-absence models and calculated the predicted area loss per population using the R package “raster”.

### Forest cover loss estimation per population

To evaluate the habitat loss risk for individual populations, we also calculated forest cover loss between 2001 and 2020 for each of the sample sites using the Global Forest Change dataset version 1.8 (Hansen et al. 2013), available at a 1 arc-second resolution (ca. 30 meters at the equator). This dataset is based on a global-scale automatized classification of Landsat 7 Enhanced Thematic Mapper Plus (ETM+) scenes. To calculate individual forest cover losses, we obtained raster values of the year of deforestation and the original forest cover bands (namely, ‘yearloss’ and ‘treecover2000’) from Hansen et al. (2013) using the Google Earth Engine version 0.1.276 (Gorelick et al. 2017) platform. We clipped these layers to each sampled population’s geographic extent using a geographic buffer of 10 km and exported the individual population rasters to R. In R, using the “raster” package, we calculated the original forest area (i.e., canopy >80%), the percentage of annual deforestation, and the variation in deforestation compared to the previous year. The future suitability projection rasters based on outlier loci were then randomly subset using the remaining forest cover since 2000 and calculated for each subpopulation to address the impacts of deforestation on the vulnerability of individual populations.

## Results

### Differences in allele frequencies between current and future environmental scenarios

From 57 individuals from six populations of *A. germinans*, we obtained 2,297 loci and identified 262 outliers with PCADapt, 154 with RDA, and 132 using both methods simultaneously. For *A. schaueriana*, we sampled 77 individuals from ten populations and recovered 6,170 loci and 224 outliers with PCADapt, 276 with RDA, and 182 with both methods simultaneously. The GF models identified the (1) annual range in air temperature, (2) precipitation seasonality, (3) mean annual air temperature, and (4) precipitation seasonality as the most important predictors for the *A. germinans* and *A. schaueriana* outliers (Table 2). The lowest values of variable importance in the gradient forest models for *A. germinans* outlier loci were the (1) annual precipitation amount, (2) growing season length, (3) number of >10 °C growing degree days, and (4) ocean surface temperature. For *A. schaueriana*, the less important predictors for outlier locus frequencies were the (1) annual precipitation amount, (2) growing season length, and (3) ocean salinity.

Future climate conditions based on differences between modern and future frequencies of putatively adaptive alleles (i.e., Procrustes differences) identified the most pronounced climate vulnerability in four *A. germinans* populations, ALC, PNB, PRC, and TMD (Fig. 3) and two *A. schaueriana* populations, ALC and VER (Fig. 4), both on the extremes of the distribution ranges of the species. Overall, the Procrustes differences were lower for *A. schaueriana* than for *A. germinans*, and the lowest mean differences for *A. germinans* were observed in the PCR and ALC populations, while those for *A. schaueriana* were observed in the FLN population.

**Fig. 3.**
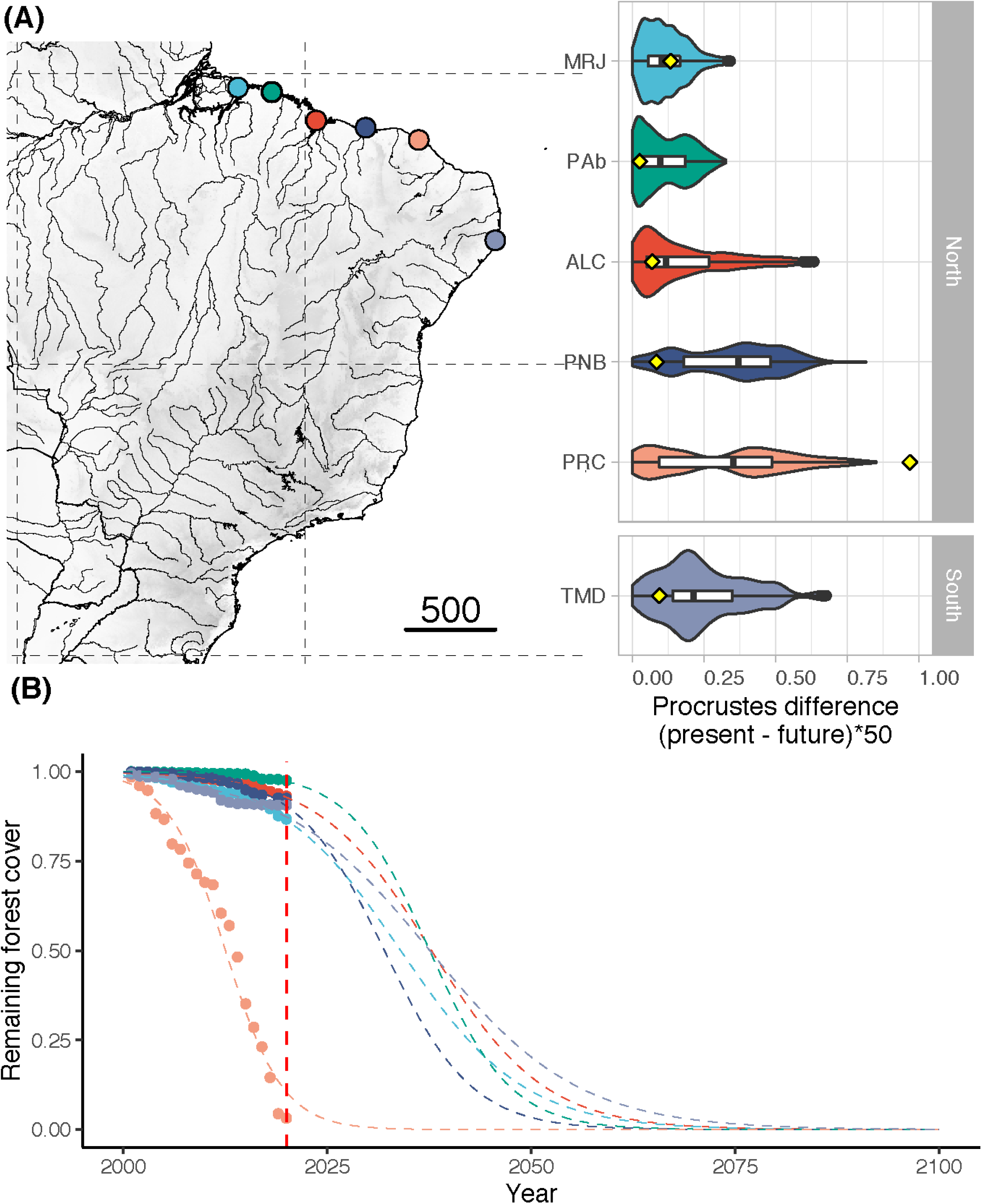
**(A)** Procrustes differences between the present and future frequencies of the outlier and reference (i.e., neutral) loci for populations of *Avicennia germinans* obtained with gradient forest and RDA/PCAdapt; the yellow dots represent the proportion of forest cover lost by 2020. **(B)** Yearly deforestation rates (2000-2020) for each *A. germinans* region included in our study.

**Fig. 4.**
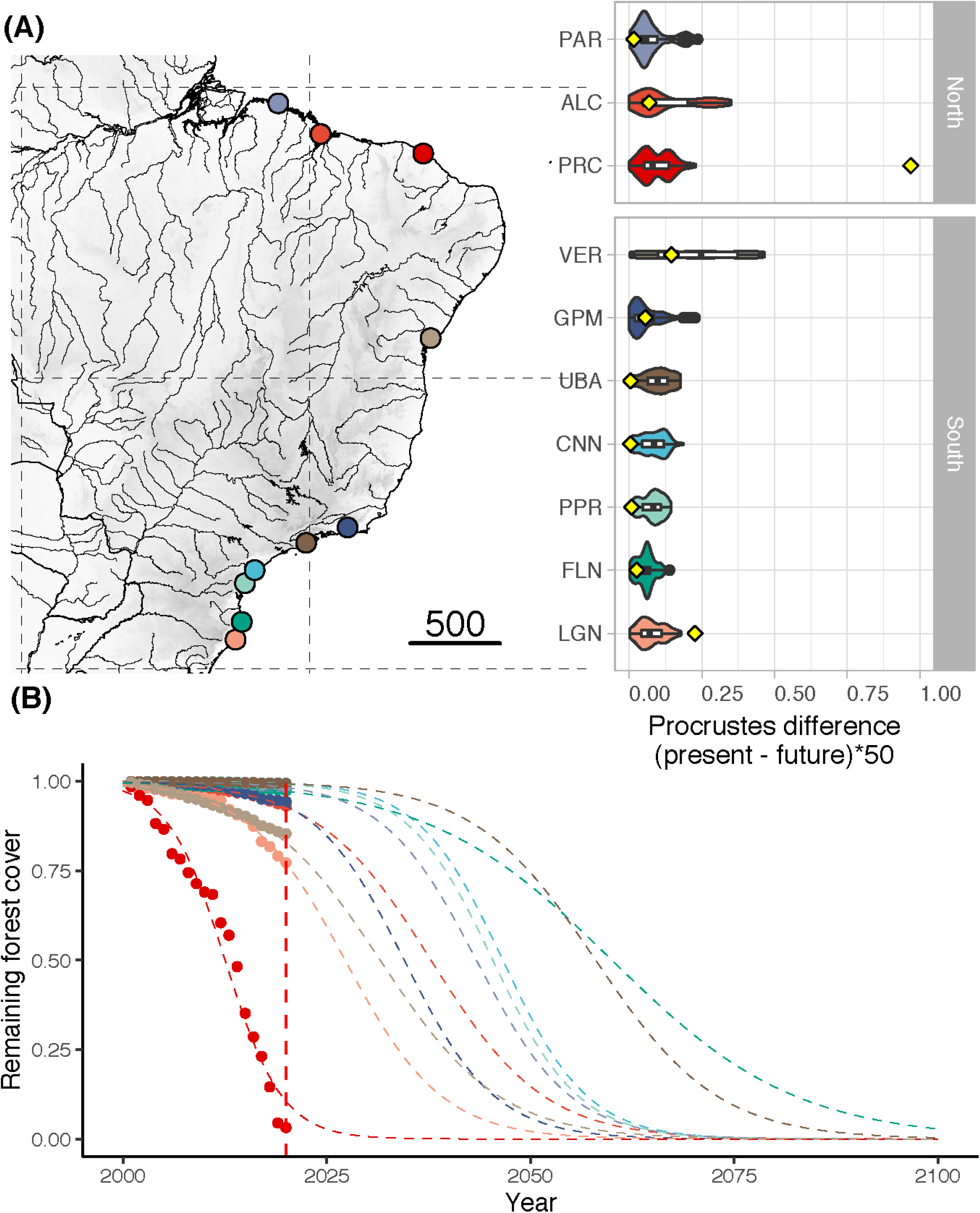
**(A)** Procrustes differences between the present and future frequencies of the outlier and reference (i.e., neutral) loci for populations of *Avicennia schaueriana* obtained with gradient forest RDA/PCAdapt; the yellow dots represent the proportion of forest cover lost by 2020. **(B)** Yearly deforestation rates (2000–2020) for each *A. schaueriana* region included in our study.

### Forest cover loss estimation per population

Since the beginning of the analyzed time series (2000), all the regions combined have lost ca. 79 km^2^ of the initial 1,350 km^2^ forest cover quantified in 2001, representing a total loss of 5.86% over two decades. The highest total forest area losses were observed in PRC and VER, with losses of 10.1 and 9.29 km^2^, respectively, accounting for 30.29% of the total observed loss. Comparatively, regions such as PPR, UBA, PAR, CNN, and PNB each lost less than 1.47 km^2^ and were the least affected by deforestation of the studied locations.

The average cover loss was substantially higher than the global rates of loss (0.4% per year) for four of the six regions with *A. germinans* (MRJ, PNB, PRC, and TMD) and a single region with *A. schaueriana* populations, namely, PRC (Table S1). Proportional cover loss per year is increasing in all the regions except TMD. Full statistical data related to the annual area loss, remaining area, loss since the previous year, loss variation, mean annual loss, and total loss are available in Supplementary Table 2. Based on the current deforestation rates, all populations show declining trends (Figures 3 and 4). *Avicennia germinans* populations present higher projected and observed deforestation rates, with PRC showing the higher forest cover loss since 2000 (96.81%). For *A. schaueriana*, the most vulnerable populations are PRC and LGN, with 96.81% and 22.72% forest cover losses since 2000, respectively.

The climate-only distribution models also showed contrasting scenarios for the north- and south-of the northeastern extremity of South America (NEESA) populations (Fig. 5). For populations of both species located at higher latitudes (TMD for *A. germinans*; UBA, PPR, CNN, and FLN for *A. schaueriana*), the models indicated an increase in the distribution range toward upper coastal areas under the SSP5-8.5 scenario by 2100. However, for populations at lower latitudes (i.e., north-NEESA), the distribution range was projected to slightly decrease. Interestingly, while both species presented an increase in total area in the southern portions of their distributions, their southernmost geographic limits did not change (remaining at 23 °S for *A. germinans* and at 28 °S for *A. schaueriana*), but a slight increase in suitability was projected for the southernmost populations, especially for *A. schaueriana*. The population-level cell count variation for each species is listed in Supplementary Table 2.

**Fig. 5.**
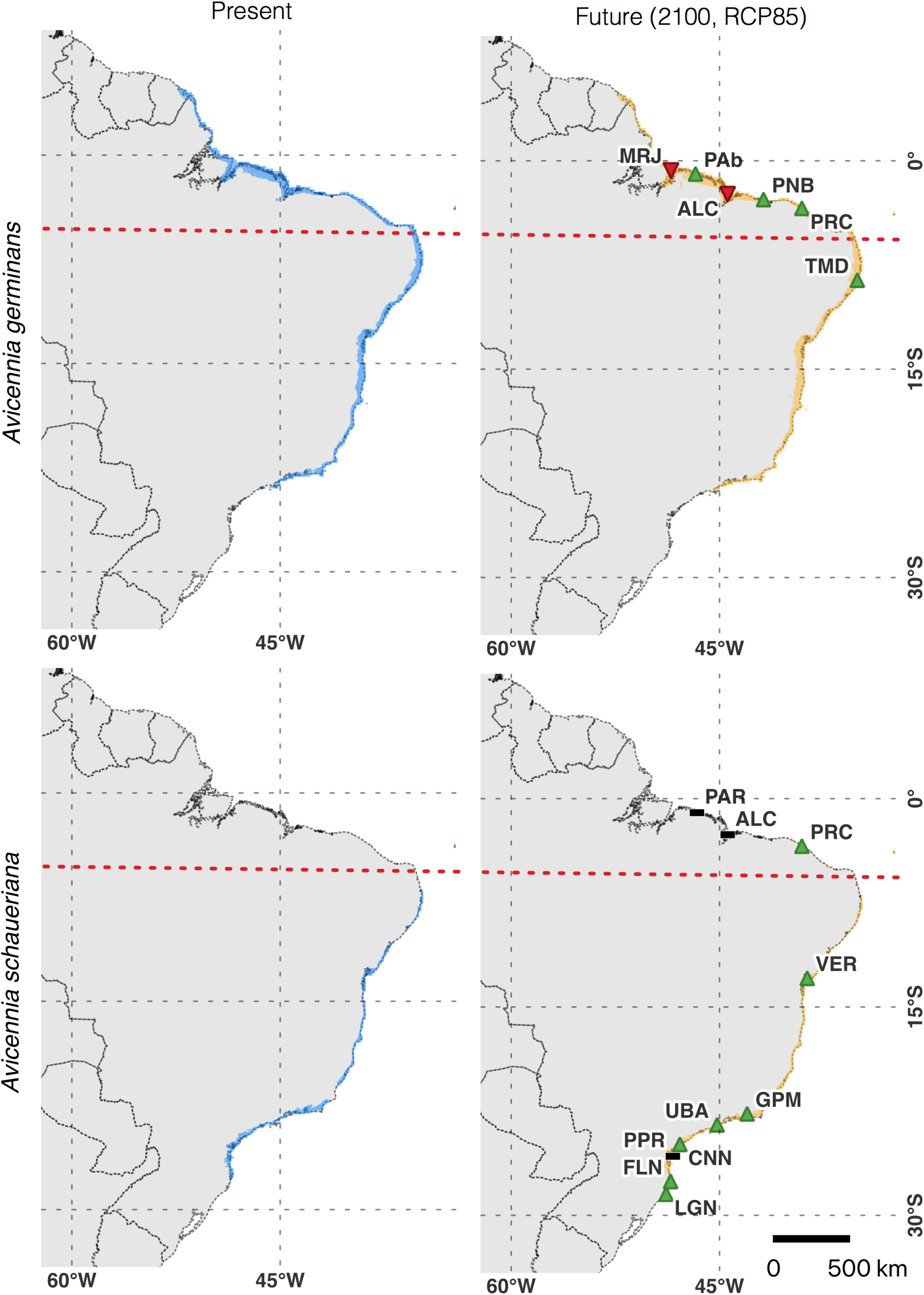
**(A)** Climate-only ensemble distribution model of *Avicennia germinans* for current and future climate scenarios and individual population trends. **(B)** Climate-only ensemble distribution model of *Avicennia schaueriana* for current and future climate scenarios and individual population trends. The dashed red line represents the north–south population division found in the northeast extremity of South America (NEESA).

## Discussion

We found evidence that *A. germinans* populations in TMD, ALC, and MRJ were the most vulnerable over the studied area based on the combined need for changes in putatively adaptive allelic frequencies and the estimated probability of climate-induced habitat loss. This result was partially consistent with our climate-only distribution models, which identified a reduced suitability for populations in ALC and MRJ but indicated an increase in suitability for populations in TMD. The recent deforestation rate was higher in PRC, MRJ, and ALC than in the other locations, with the PRC region having lost a remarkably high fraction (28%) of forest cover area from 2019 to 2020. For *A. schaueriana*, the VER population showed a higher genome-environmental vulnerability, and the ALC, GPM, and LGN populations showed higher deforestation percentages.

The climate-only models showed a slight decrease in suitability for two equatorial populations (MRJ and ALC), while other subtropical areas (with the exception of PPR) were projected to increase in suitability. Our models suggest that the rates of deforestation among populations in subtropical areas are higher than the global average rates of deforestation. These results may indicate concerning scenarios for subtropical populations due to the effects of the dangerous combination of limited gene flow, habitat fragmentation, connectivity loss, climate-induced habitat loss, and limited population gene pools on the long-term persistence of Brazilian mangroves. The projected forest cover loss for the *A. germinans* populations PRC and TMD indicates that some populations may be more vulnerable to land-use change and deforestation than to the potential risks of climate-induced habitat loss. Both of these areas also present high values of Procrustes differences for the frequencies of potentially selective loci, ranking them as the most vulnerable populations in our analysis.

### Differences in the vulnerability of individual populations to climatic change

Overall, the mean Procrustes differences were not significantly different between the northern and southern populations of either species. However, higher individual Procrustes differences—i.e., higher counts of grid cells requiring the greatest changes in allele frequencies—were observed at intermediate latitudes in the PNB, PRC, and TMD populations of *A. germinans* and in the VER population of *A. schaueriana*. Furthermore, for *A. schaueriana*, the population vulnerability estimated by the mean Procrustes differences was lower toward the southern and northern areas relative to the VER population. As higher mismatches in allele frequencies were calculated for populations located at intermediate latitudes, we presume that alleles putatively associated with better fitness under future conditions are currently found at low frequencies at these sites.

We define the northern and southern regions of NEESA as core groups related to genomic diversity. The PNB, PRC, TMD and VER populations occupy peripheral positions in relation to the range of both species (Fig. 2) and could be subjected to higher risks of extinction due to these peripheral environments (Hardie & Hutchings, 2010). The results identified by our models support the findings of Cruz et al. (2019) and Cruz et al. (2020), who identified contrasting allelic frequencies in the genes linked to decisive environmental pressures in central and marginal *Avicennia* populations in Brazil. Populations north of NEESA are mostly linked to equatorial climate and have alleles that are more frequently specifically linked to genes involved in regulation and response to light and saline saturation; in contrast, populations farther south along the Brazilian coast have alleles that are more linked to response to cold and water balance (Cruz et al. 2019, Cruz et al. 2020). In comparison, marginal subtropical and marginal equatorial populations present lower genetic diversities for these loci and reduced population sizes in comparison with the stable core populations.

The identification of this nonlinear response of loci to environmental gradients emphasizes the role of dispersal and gene flow between populations as a fundamental condition to ensure effective conservation of these forests, especially at intermediate latitudes. By sharing alleles through seedlings and pollen, populations across the geographic distribution of a species will be better suited to cope with novel environmental pressures due to increases in adaptive resources. However, since Brazilian mangroves are partially isolated in two geographic clusters (i.e., south-NEESA and north-NEESA), it is reasonable to assume that alleles with higher adaptive values may not be sufficiently shared between these populations through propagule dispersion. Thus, enhancing connectivity between the remaining fragments of mangrove forests, while necessary to ensure gene flow, might not be sufficient to provide the genomic resources necessary to withstand the warmer and drier conditions predicted for the next century. Therefore, based on the evidence reported here, we suggest that, in addition to enhancing connectivity between fragments, artificially inducing gene flow from north to south NEESA should be considered in the genetic management plans of *A. germinans* and *A. schaueriana*. This method has been deemed successful and recommended by Kottler et al. (2021) and Lidelli et al. (2021). Nonetheless, the allelic composition of the diversity sources must be carefully chosen to promote allelic combinations that may confer local adaptations to the transitional environmental conditions to which TMD and VER populations might be exposed (Cruz et al. 2019, Cruz et al. 2020, Silva-Pereira et al. 2020). The benefits of this approach will likely overcome the risks associated with separate management of north- and south-NEESA populations, as suggested in various studies discussing conservation approaches for isolated populations (Frankham 2015, Liddell et al. 2021).

The results obtained for the identification of outlier loci, which were the foundation of the modeling step, require careful interpretation. Many factors may influence the results of selective locus identification, and sample size (especially if the number of loci is low) could inflate the detection of false-positive results by the algorithm used to identify the loci under selection (PCAdapt and RDA). Therefore, our results and the interpretation presented here are in the context of previous results presented for this dataset (i.e., Cruz et al., 2019, Silva et al., 2021), which support the strong influence of climate and precipitation on the local adaptation of *Avicennia* populations in Brazil.

### Future climate scenarios reinforce abiotic stress in mangroves on the Brazilian coast

Future projections of climatic variables indicate general warming and aridification trends for the Brazilian coast (Fig. 2). Surprisingly, niche models for *A. germinans* and *A. schaueriana* indicate an overall stability or increase in distribution ranges, with very slight reductions in environmentally suitable areas only for MRJ and ALC, two northern populations of *A. germinans* (Fig. 5). As previously reported (Mori et al. 2015, Bajay et al. 2018, Cruz et al. 2019 and 2020) and demonstrated in this study, the equatorial and subtropical portions of the Brazilian coast consist of distinct adaptive and demographic groups that will face different outcomes in a warmer and drier climate. Compared to subtropical latitudes, the equatorial region where *A. germinans* and *A. schaueriana* co-occur is hotter and drier (Fig. 2). Plants in this region show signs of heat stress in the field, with high expression levels of the heat shock protein-coding genes HSP17, 70, and 101 and the transcription factor RAP2.3 (ethylene-responsive binding factor) (Cruz et al. 2019). Similarly, for another dominant mangrove tree, *Rhizophora mangle* (*Rhizophoraceae*), equatorial populations were shown to be under severe stress caused by excessive heat (Bajay et al. 2018). Such evidence indicates that the increase in temperature in these regions will severely impact the mangrove tree populations, which are already under thermal stress.

The subtropical populations, however, will likely face different challenges. First, the increase in annual temperature can reduce the occurrence of freezing events, which are a key limiting factor for the occurrence of mangroves at high latitudes (Cook-Patton et al. 2015, Osland et al. 2020). Second, the reduction in precipitation combined with warming will lead to an increased atmospheric water vapor pressure deficit, which can have harmful effects on mangrove populations. Cruz et al. (2019) reported potential local adaptations that confer hydraulic and transpiration systems suitable for higher water availability on subtropical *A. schaueriana* populations, making them more susceptible to desiccation and cavitation than populations from warmer and drier equatorial regions (Markestejin et al. 2011).

For *A. germinans*, genetic variation with evidence of selection correlates with precipitation regimens and, more specifically, with the combination of the driest and coldest quarters of the year. These results corroborate those reported by Silva et al. (2021), who identified environmental isolation with patterns along atmospheric temperature, precipitation, and solar radiation gradients as the model that best explained genetic differentiation between populations of this species, and Cruz et al. (2020), who found that the molecular responses of *A. germinans* populations in these localities were associated with freshwater limitations and were more remarkable north of the South Equatorial Current (SEC), more precisely, in sites where freshwater inflows by rivers are scarcer. Precipitation regimes have commonly been related to coastal wetland forest distributions (Cavanaugh et al. 2018, Osland et al. 2015, Osland et al. 2016).

Freshwater variation also seems to be an important factor determining the vulnerability of populations of *A. germinans* to the climatic conditions predicted for the end of this century. We found greater future mean genetic displacement (Procrustes differences) in the TMD populations, followed by the MRJ and ALC populations. These populations currently face lower hydraulic stress due to a greater amplitude in the rainfall regime (TMD) and a greater inflow of freshwater from Amazonian rivers (MRJ). However, according to the climate predictions, they may be more likely to experience higher temperatures and drier climates and, consequently, increased water salinity. In addition, for the TMD population—and probably for the populations existing at the distribution limit of this species (ca. 22°S)—the bifurcation caused by the SEC and the north–south direction of the Brazilian current (BC) further limit the gene flow and genetic input that this population will need to withstand drier and hotter climates.

According to the genotype-environment association analyses, variations in temperature and precipitation patterns were important factors determining the genetic differentiation of *A. schaueriana* along the Brazilian coast. These results were consistent with those obtained by Cruz et al. (2019), who found variations in the loci present in the genomic regions functionally associated with biological processes related to responses to temperature, solar radiation, and freshwater availability, such as the response to osmotic stress, anthocyanin biosynthesis, protection against ultraviolet rays (UV), and biogenesis of the components of the photosynthetic apparatus. Silva et al. (2021) showed that for *A. schaueriana*, the environmental gradients of temperature and precipitation were closely correlated with the geographic variations represented by the latitudinal gradient.

We identified pronounced Procrustes differences between the frequencies of present and future outlier loci for the VER population (Veracruz-BA). This population is found south of the NEESA in a region with a lower average rainfall and higher temperatures than regions further south along the coast, such as UBA, CNN, PPR, FLN, and LGN. However, the variation in the future precipitation scenarios for VER is greater than that for all the other locations with populations of *A. schaueriana* (Fig. 2), suggesting drought as a potential stressor in the future. This scenario is also aggravated by the limited movement of propagules from the populations further south to the population in VER; the north–south direction of the BC restricts dispersal, limiting the exchange of genetic material between populations to the north and south of its bifurcation. For these localities, we recommend that reforestation measures should include seedlings with a higher tolerance to drought as well as local specimens.

### Increase in deforestation rate in the last two decades

We quantified alarmingly high and increasing rates of vegetation cover loss for Brazilian mangrove forests, with 15 out of 16 sites showing increasing trends and 10 out of 16 sites showing yearly percentage losses higher than the average global loss rate. A high recent deforestation rate was observed in the populations in PRC, MRJ, and ALC. Combined with the predicted future genomic-environmental pressures, these cover losses decrease the long-term resilience of these populations and threaten the maintenance of ecosystem services essential to mangrove-associated species and the human populations that rely on them.

Deforestation in mangroves affects multiple ecosystem services, but perhaps the most important consequence is the disproportionate loss in organic carbon fixation per area. It is estimated that Brazil accounts for more than 9% of the global mangrove-based carbon stock in its 7,674 km^2^ area of mangrove forests (Hamilton & Friess, 2018); thus, land-use changes in this broad area can become significant sources of carbon emissions when the forests are removed. Locally, deforestation can lead to the disruption of ecological processes, an increase in edge effects, and the loss of multiple levels of genetic diversity, reducing the capacity of local communities to adapt to environmental changes (Baucom et al. 2005, Haddad et al. 2015). Available studies assessing the remaining genetic diversity of other mangrove species such as *Rhizophora apiculata*, a mangrove tree from the Eastern Hemisphere, identified high percentages of homozygotes in their populations, suggesting persistent inbreeding, which was attributed to habitat fragmentation and persistent low population sizes caused by deforestation (Azman et al. 2020). Low effective population sizes were also reported for mangroves in other regions of Asia, such as the Indo-Malayan coast, where lower genetic diversity greatly increased the vulnerability of less genetically diverse mangrove species to coastal flooding and sea-level variations (Guo et al. 2018). Recently, a global mangrove deforestation survey (Bryan-Brown et al. 2020) reported Brazilian mangroves to be hotspots of habitat loss, yet with “*lower rates of fragmentation*” compared to the fragmentation rates in other countries assessed in their study. However, this scenario is likely to change due to the rapid pace of forest cover loss that we identified. Our results indicate that mangrove population sizes and connectivity are likely to decrease, thus, deforestation may be an even more urgent threat to Brazilian mangroves than climate change because it is deteriorating the already limited evolutionary potential of its populations.

### Poleward migration in A. schaueriana and A. germinans

The potential for range expansion toward higher latitudes was not identified by our ensemble distribution models. However, the higher vulnerability of the intermediate-latitude populations identified by the combined genomic-environmental models supports this scenario, which has been hypothesized for Brazil (Soares et al. 2012) and reported for regions on the west coast of South America (Saintlan et al. 2014) and in other places in the world, such as North America (Osland et al. 2020), South Africa, Australia, and Asia (Saintlan et al. 2014). Globally, however, satellite imagery and literature reviews showed no evidence that mangroves are undergoing unassisted distribution shifts to higher latitudes, even with temperatures increasing in their current distributions (Hickey et al. 2017). Therefore, Brazilian mangroves may be ‘trapped’ inside their current latitudinal distribution area, with their potential to adapt to the warming and drying conditions in this area strongly decreased by habitat loss and low connectivity. In Brazil, the southern limit of mangrove forests is located in Santo Antonio Lagoon in the municipality of Laguna (28°28’S; 48°50’W) (Soares et al. 2012), and this southern limit has not changed since at least 1990 (Saintlan et al. 2014). This geographic limitation is, however, most likely attributed to a restricted dispersal due to a local ocean current rather than to the environmental suitability of the region (Soares et al. 2012, Saintlan et al. 2014).

Given the environmental factors to which the studied species and populations are most sensitive, we recommend that conservation measures must take into account the adaptive particularities of each population along the Brazilian coast. Mitigating actions should be aimed not only at increasing connectivity and reducing fragmentation, as mentioned in the previous sections, but also at comprehensively sampling the functional factors of effective ecological restoration actions. Surprisingly, despite harboring a considerable fraction of global mangrove forests, South American mangroves are considerably underrepresented in the literature compared to Southeast Asian or Australian mangroves (e.g., Gorman 2018). Site-based conservation is essential because it allows the long-term persistence of many species by sustaining viable populations in their natural states. Our results demonstrate one of several potential applications of community-level modeling of genomic variation to improve predictions of the effects of climate change on population-level vulnerability, which is an important advancement in environmental-genomic biodiversity models.

## Contribution to the field

Our research evaluates the vulnerability of two species of Brazilian mangrove trees to climate-induced habitat loss using a genomic-environmental approach. We were able to demonstrate that populations found at intermediate latitudes along the Brazilian coastal gradient will be more vulnerable to higher temperatures and drier conditions than clearly equatorial or subtropical populations of either species, mostly due to local adaptations. We also quantified recent forest cover loss for each area and predicted the loss of suitability due to climate only to provide robust estimates of vulnerability for each population. Our results provide an important resource for conservation planning and demonstrate the potential application of novel genomic-environmental methods.

## Ethics statements

### Studies involving animal subjects

Generated Statement: No animal studies are presented in this manuscript.

### Studies involving human subjects

Generated Statement: No human studies are presented in this manuscript.

### Inclusion of identifiable human data

Generated Statement: No potentially identifiable human images or data is presented in this study.

### Data availability statement

Generated Statement: The original contributions presented in the study are included in the article/supplementary material, further inquiries can be directed to the corresponding author/s.

## Supplementary material

**Supplementary Figure 1.** Manhattan plots for **(A)** *Avicennia schaueriana* and **(B)** *Avicennia germinans* loci identified by PCAdapt as SNPs with MAF > 0.05.

**Supplementary Figure 2.** Score plots of the sampled individuals of **(A)** *Avicennia germinans* and **(B)** *Avicennia schaueriana* and the axis (PC1 and PC2) with the greatest explanatory power for population structure (identified by PCAdapt).

**Supplementary Figure 3.** Total forest cover loss (in square kilometers) during 2001–2020 for each mangrove region (10 km buffer) sampled in this study. The data were calculated using the Global Forest Change dataset version 1.8 (Hansen et al. 2013).

**Supplementary Table 1.** Geographic and environmental information on the *Avicennia germinans* and *Avicennia schaueriana* populations sampled in this study.

**Supplementary Table 2.** Forest cover losses and climate suitability changes for the *Avicennia germinans* and *Avicennia schaueriana* populations sampled in this study.

**Supplementary Table 3.** *Avicennia germinans* and *Avicennia schaueriana* occurrence records used for the modeling step.

**Supplementary Data 1.** R script used for generating the genomic-environmental models.

## 2. Author Contributions

JDV developed the methodology, prepared the figures, and wrote the foundational manuscript and supplementary data. GMM conceived of the study, provided data and assisted with writing. MVC conceived the study, generated the genomic data and analyzed the data. MFS assisted with the Bayenv2 analysis, discussion of the results and manuscript writing. YAM assisted with the manuscript writing and discussion of the results. APS conceived of the study and provided project leadership.

## 4. Acknowledgments

JDV thanks Dr. Paul Andrew Maier for sharing a brilliant tutorial on how to implement GF models in genomic-environmental association studies.

## 5. Data Availability Statement

The datasets are available as supplementary material.

